# Dissociable Effects of Emotional Stimuli on Perception and Decision-Making for Time

**DOI:** 10.1101/2020.04.24.059717

**Authors:** Keri Gladhill, Giovanna Mioni, Martin Wiener

## Abstract

Previous research has demonstrated that negative emotional faces dilate time perception, however, the mechanisms underlying this phenomenon are not fully understood. Previous attempts focus on the pacemaker-accumulator model of time perception, which includes a clock, memory, and decision-making stage, wherein emotion affects one of these stages; possibly by increasing pacemaker rate via arousal, increasing accumulation rate via attention, or by biasing decision-making. To further investigate the stage(s) that emotion is affecting time perception we conducted a visual temporal bisection task with sub-second intervals while recording 64-channel electroencephalogram (EEG). To separate the influence of face and timing responses the temporal stimulus was preceded and followed by a face stimulus displaying a neutral or negative expression creating three trial-types: Neg→Neut, Neut→Neg, or Neut→Neut. The data revealed a leftward shift in bisection point (BP) in Neg→Neut and Neut→Neg suggesting an overestimation of time. Neurally, we found the face-responsive N170 component was larger for negative faces and the N1 and contingent negative variation (CNV) were larger when preceded by a negative face. We also found an interaction effect between condition and response for the late positive component of timing (LPCt) and a significant difference between response (short/long) in the neutral condition. We conclude that a preceding negative face affects the clock stage leading to more pulses being accumulated, either through attention or arousal, as indexed by a larger N1, CNV, and N170; whereas viewing the negative face second biased decision-making leading to “short” responses being less likely, as evidenced by the LPCt.

## Introduction

The ability to perceive time accurately is invaluable to humans. It is important in everyday activities to understand when in time and for how long an event occurred, or how soon in the future an event might occur. These different aspects of time allow us to move through the world and perform necessary actions on time. It is necessary, then, that we understand the conditions under which time perception can be altered and the inaccuracies that result. Emotion and the passing of time are closely linked in the human language; for example, we often comment on time passing more quickly when having fun and time passing more slowly during distressing events. Given this close link, it is not surprising that emotion has been found to alter one’s perception of time (Lake et al., 2016). Multiple theories have been presented to explain how emotion alters time perception; the focus of the current study is to tease apart differing explanations between them and compare contributing factors.

The pacemaker-accumulator model of time perception, or scalar expectancy theory (SET), first postulated by Treisman (1963), and later expanded by Gibbon, Church, and Meck (1984), includes three main stages: the clock, memory, and decision-making (Figure 1). The clock stage involves a pacemaker which constantly produces pulses; at the onset of a to-be-timed event, these pulses are then collected by an accumulator at a rate that is modulated by a switch- or gate-like mechanism (Zakay & Block, 1995). The memory stage involves comparing the duration of an interval based on the number of pulses collected in working memory to previously stored values in long-term memory. Last is the decision-making stage which involves making a judgment regarding the previously made comparison. This decision can be made by comparing a temporal stimulus with another of a different duration, by reproducing a time interval, or by determining when to perform an action (Allman et al., 2014, Lake et al., 2016).

**Figure 1.**
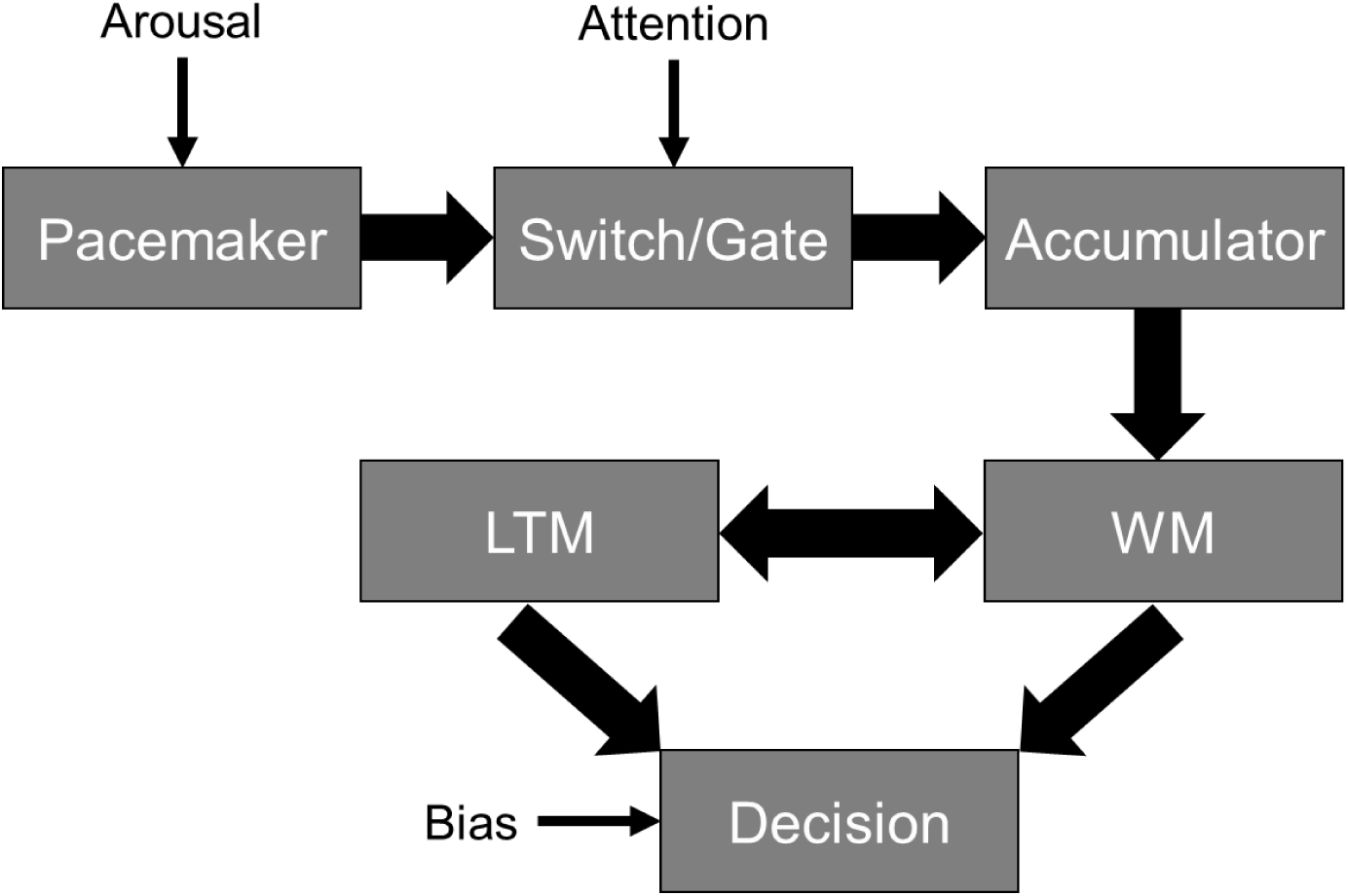
Pacemaker-accumulator model of time perception/scalar expectancy theory (SET) includes three main stages: the clock, memory, and decision-making. Emotion may affect arousal via the pacemaker, attention via the switch/gate, and/or bias the decision.

Emotion has been proposed to impact time by acting on distinct stages in the pacemaker accumulator model. First, arousal resulting from emotional stimuli can increase the rate of the pacemaker, which in turn leads to a longer perception of time. For example, the emotional states that involve higher levels of arousal (e.g. anger or fear) would increase the pacemaker rate more compared to emotions with lower levels of arousal (e.g. sad). However, emotions with lower levels of arousal would still increase the rate of the pacemaker beyond that of neutral emotions (Droit-Volet, et al. 2004).

Second, changes in attention due to emotional stimuli can alter the distribution of cognitive resources to, or away from, a timing task thus affecting the gate or switch function of the model (Lake et al., 2016). Lui and colleagues (2011) suggested that an emotional temporal stimulus will increase attention causing the switch to flicker closed less which will leads to a larger number of pulses accumulated thus leading to an overestimation; whereas, emotional stimuli presented before the temporal stimuli will cause an allocation of resources away from the subsequent temporal stimuli causing the switch to flicker closed more which leads to a smaller number of pulses accumulated thus leading to an underestimation. However, discerning between arousal or attentional affects remains difficult, because both can lead to identical behavioral effects (Burle & Casini, 2001).

Third, emotion could be causing a bias at the decision-making stage of the model. Previous research by Lieving and colleagues (2006), has shown that an increased delay between sample stimuli and choice results in a “choose-short” effect. In this situation it is thought that subjects exhibit a bias towards the shorter choice when the retention interval is long because their memory for the stimulus duration is decaying as the delay increases. In Lieving and colleagues’ study they varied the retention interval (RI) by 0, 8, 16, and 32 seconds between and within sessions and found a right-ward shift in the bisection point (BP) for each of the delays compared to the no-delay baseline which suggests that participants experienced a bias to choose short.

Research on emotional timing has utilized emotional faces, words, and sounds as well as other emotional stimuli from the International Affective Picture System (IAPS). Previous research has found that emotional faces cause an overestimation of time perception; specifically, it has been shown that angry faces elicit the largest overestimation compared to happy, sad, or neutral expressions with no differences in happy or sad expressions (Droit-Volet et al., 2004; Fayolle & Droit-Volet, 2014; Tipples et al., 2015). In addition to behavioral findings, we examined electrophysiological data using electroencephalography (EEG) recordings; specifically, we analyzed event-related potentials (ERPs) that have been associated with timing in the context of emotional stimuli.

The contingent negative variation (CNV) is strongly associated with time perception and is thought to be an index of timing, with the amplitude being found to increase in relation to the length of the temporal estimates. This ERP was first described by Walter and colleagues (1964) as a slow negative wave that occurs between a warning and imperative stimulus. In addition, preceding the CNV, a brief negative/positive deflection is often observed (N1/P2), with the N1 component thought to indicate selective attention where a more negative amplitude indicates increased attention (Brunia, et al., 2011). Numerous studies have now demonstrated an association between both the N1 and CNV during time estimation (Pouthas et al., 2000; Xuan et al., 2009; Wiener et al., 2012).

The N170/vertex positive potential (VPP) are ERPs associated with face perception and emotion. The VPP is a large positive potential that peaks at the vertex between 140-180ms after the onset of face stimuli (Jeffreys et al., 1989). In contrast, the N170 corresponds to the visual N1 component which is the first negative deflection on posterior scalp regions (Bentin, et al. 1996). The N170 has the earliest and strongest difference in amplitude between faces and non-faces with the magnitude being larger for emotional faces, specifically fearful and angry, compared to neutral faces (Frühholz et al., 2011; Hajcak et al., 2011).

The late positive component of timing (LPCt) is an additional timing-related ERP thought to be associated with decision-making and difficulty in temporal discrimination (Paul et al., 2003, 2011; Gontier et al., 2009; Wiener & Thompson, 2015), which appears as a positive frontocentral deflection following the CNV. The time point in which the LPCt occurs in these cases varies depending on the amount of time until response with some studies finding the LPCt at stimulus offset and others immediately following the subject response (i.e. button press). The aforementioned studies have also found the LPCt to covary with temporal duration and Wiener and Thompson (2015) found the LPCt to vary based on choice (short or long).

While previous evidence suggests emotional stimuli cause an overestimation in time perception and that there are clear ERPs that are elicited when timing and viewing emotional stimuli; it is not yet understood whether emotion is affecting the clock portion of the pacemaker accumulator model via arousal and/or attention or the decision-making portion due to bias. The purpose of this study is an attempt to differentiate between these two possibilities by combining a temporal bisection task with ERP analysis. Whereas previous ERP studies of emotional effects on time perception had subjects measure the duration of emotional stimuli directly, the innovation of the current study is that the stimulus to be timed is not an emotional stimulus and, unlike Lui and colleagues (2011), was temporally flanked by emotional stimuli, presented before and after the timed stimulus, with one stimulus having an emotional valence (negative) and the other not (neutral). The use of this design was twofold: first, when timing an emotional stimulus, the resultant changes in time perception may arise from either clock-mediated or decision-mediated effects, as described above, but are intermixed during the presentation of the emotional stimulus. By flanking the timed stimulus with an emotional stimulus before or after the timed stimulus, it is therefore possible to determine which stage(s) emotion is affecting. Second, in previous ERP studies of emotion and time perception (Gan, et al. 2009; Tamm, et al. 2014; Zhang, et al. 2014), the timing-related ERP components (i.e.N1, CNV) will similarly be intermixed with emotion or face-processing components (i.e. N170, VPP). As such, by temporally separating the emotional stimuli from the timed ones, we could account for any independent effects or contributing effects from either response.

We hypothesized that (1) if emotion was only affecting the clock portion of the model that participants would overestimate the duration when a negative face was shown *before* the temporal stimulus but (2) if emotion was only affecting the decision-making portion of the model, that participants would overestimate the duration when a negative face was shown *after* the temporal stimulus. Correspondingly, if emotion was affecting the clock-stage, then an increase in the CNV would be expected when an emotional stimulus precedes the temporal stimulus (Neg→Neut); whereas if emotion impacted the decision-stage, then the response-locked LPCt should be affected when an emotional stimulus follows the temporal stimulus (Neut→Neg). More specifically, if the first hypothesis is supported, we also hypothesize that (1a) if it is arousal that is affecting the clock, we should expect a more negative N170 amplitude when viewing the negative face compared to viewing the neutral face; whereas (1b) if it is attention causing the affect we should expect a more negative N1 amplitude when viewing the negative face compared to viewing the neutral face.

## Method

### Participants

This study included 19 right-handed participants (13 females, 4 males, 2 undisclosed; 18-27 years old) from the undergraduate student population of George Mason University. Two participants were removed as outliers (one for EEG data and one for behavioral performance). All participants had normal or corrected-to-normal vision. Participants were compensated for their time with either research credits or monetary payment. All participants completed a demographic questionnaire and provided informed consent as approved by The Institutional Review Board at George Mason University.

### Apparatus and procedure

Participants completed a temporal bisection task on a computer in which they categorized the duration of a temporal stimulus as either short or long by pressing ‘S’ or ‘L’ on the keyboard, respectively. Prior to and following the temporal stimulus, participants saw a human face which expressed either a negative emotion (anger) or a neutral emotion. The face before and after the stimuli was always the same individual and the combinations of faces presented were randomly selected from four male and four female faces from trial to trial. Each trial was classified into one of three conditions: negative first/neutral second (Neg→Neut), neutral first/negative second (Neut→Neg), or neutral first/neutral second (Neut→Neut). The study consisted of three blocks with 168 trials in each. Each trial proceeded as follows: a fixation cross for 1000ms, the first face for 600ms, a 500ms blank screen, the temporal stimulus for one of seven log-spaced durations (300, 360, 433, 520, 624, 749, or 900ms), a 500ms blank screen, the second face for 600ms, a 500ms blank screen, and the word “Respond” until the participant responded by pressing either ‘S’ or ‘L’ (Figure 2). Each trial lasted approximately 4-4.6 seconds and all three blocks took approximately 50-55 minutes.

**Figure 2.**
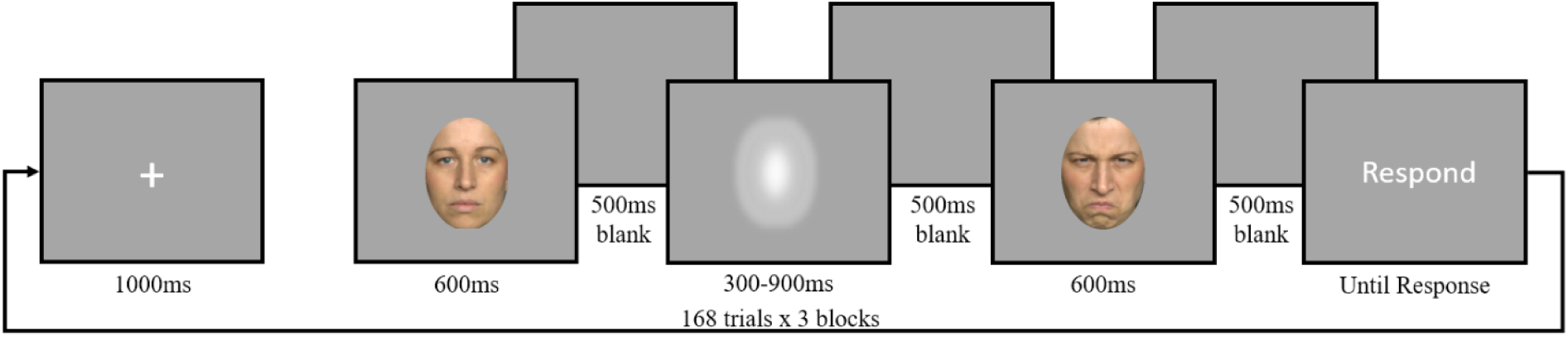
Study paradigm: fixation cross (1000ms), face stimulus (600ms), blank screen (500ms), Gaussian blur (300, 360, 433, 520, 624, 749, or 900ms), blank screen (500ms), face stimulus (600ms), blank screen (500ms), “Respond” (until key press). Example of Neut→Neg trial.

Participants were seated approximately 36 inches in front of a 31.5” LED monitor built by Cambridge Research Systems (120 Hz refresh rate). The visual stimuli were presented using PsychoPy2 (v1.90.3; Pierce, 2007). The emotional stimuli included pictures of four different male and four different female faces showing an angry and a neutral expression selected from the FACES dataset (Ebner et al., 2010; Images: 008_y_m_a_b, 008_y_m_n_b, 066_y_m_a_b, 06_y_m_n_b), 010_y_f_a_b, 010_y_f_n_b, 040_y_f_a_b, 040_y_f_n_b). The temporal stimulus was a Gaussian blur presented at the same size and location as the emotional stimuli. Participants were given instructions on the task both verbally and written and were instructed to minimize head and eye movements during the trials.

### Data collection

The experimental software (PsychoPy2) recorded response time (time from when “Respond” appeared on the screen until the button press) and key presses (‘S’ or ‘L’). EEG data was recorded from 64-actiCAP Slim electrodes connected to an actiCHamp amplifier (Brain Products) using BrainVision Recorder software. The EEG data were sampled at 1000 Hz.

### Data analysis

We analyzed the bisection point (BP) in each of the possible conditions across the different temporal durations. The BP is derived from the proportion of long responses. Specifically, behavioral data were first analyzed in terms of the proportion of “long” responses of each participant for each of the temporal intervals and for each of the emotional conditions. From this measurement, a psychometric function was fit to each individual subject’s data. Each psychometric function consisted of a cumulative Gaussian function fit via maximum likelihood estimation procedure. From these fits, we then extracted the BP and the standard deviation (σ), which is an estimate of the difference threshold. The BP is an index of perceived duration and can be described as the time value corresponding to the 0.5 probability of “long” responses. We also calculated the Coefficient of Variation (CV) as the standard deviation divided by the BP with higher CVs corresponding to lower temporal sensitivity. Repeated measures ANOVAs were conducted on BP, CV, and RT (reaction time) across conditions (Neg→Neut, Neut→Neg, Neut→Neut) as within-subject factor. RT was also analyzed across temporal intervals.

EEG data were down sampled from 1000 Hz to 500 Hz and re-referenced to the average of the mastoid channels (T9 and T10) and then epoched from 500ms before to 1000ms after the facial stimuli onset, temporal stimulus onset, and the key press response onset. Independent Component Analysis (ICA) was employed to detect and remove artifacts related to eye blinks, muscle, and line noise artifacts, after which a 1-50 Hz finite impulse response bandpass filter was applied. Separate epochs were generated for the face stimuli for each of the four face conditions (negative first, neutral first, negative second, neutral second), for the temporal stimulus for each face combination condition, collapsed across duration (Neg→Neut, Neut→Neg, Neut→Neut), response (short: S, long: L), and condition (Neg→Neut_S, Neg→Neut_L, Neut→Neg_S, Neut→Neg_L, Neut→Neut_S, Neut→Neut_L).

In order to analyze the N170 we calculated the mean amplitude across participants for negative faces and neutral faces at parietal and parieto-occipital electrodes (P7, P8, PO7, PO8) between 145-185ms (Nemrodov et al., 2016). For the N1 and CNV, we analyzed the mean amplitude within a frontocentral a-priori cluster of nine electrodes centered on FCz (FCz, Fz, Cz, F1, F2, FC2, C1, C2), within two time windows: 150-190ms and 250-450ms, respectively (Wiener et al. 2012; Wiener & Thompson, 2015). The mean amplitude of the LPCt was analyzed between 200-600ms after the response at the same nine electrodes as the CNV and N1 (Wiener & Thompson, 2015).

## Results

### Behavioral effects

Results showed no significant effect of condition on the BP (*F*_(2,36)_ = 1.22, *p* = 0.31; Figure 3). Although not significant, we observed a minor leftward shift in the BP with a marginally larger shift for Neg→Neut and Neut→Neg, suggesting a slight overestimation of time perception in those conditions (BP – Neut→Neut: M = 590.00, SE = 17.54, 95% CI = [555.83, 624.57]; Neut→Neg: M = 579.00, SE = 16.79, 95% CI = [546.10, 611.90]; Neg→Neut: M = 578.00, SE = 17.04, 95% CI = [544.59, 611.41]). A repeated measures ANOVA was also conducted for CV and for reaction time (RT), two participants were not included in the RT analyses due to missing data. Neither analysis revealed significant results (*F*_(2,36)_ = 0.02, *p* = 0.98; *F*_(2,32)_ = 0.33, *p* = 0.72, respectively). CV – Neut→Neut: M = 0.32, SE = 0.03, 95% CI =[0.25, 0.39]; Neut→Neg: M = 0.32, SE =0.03, 95% CI = [0.25, 0.39]; Neg→Neut: M = 0.32, SE = 0.04, 95% CI = [0.25, 0.39]; RT – Neut→Neut: M = 600.40, SE = 59.29, 95% CI =[490, 710]; Neut→Neg: M = 592.60, SE = 59.74, 95% CI = [476, 710]; Neg→Neut: M = 608.00, SE= 58.55, 95% CI = [493, 723]).

**Figure 3.**
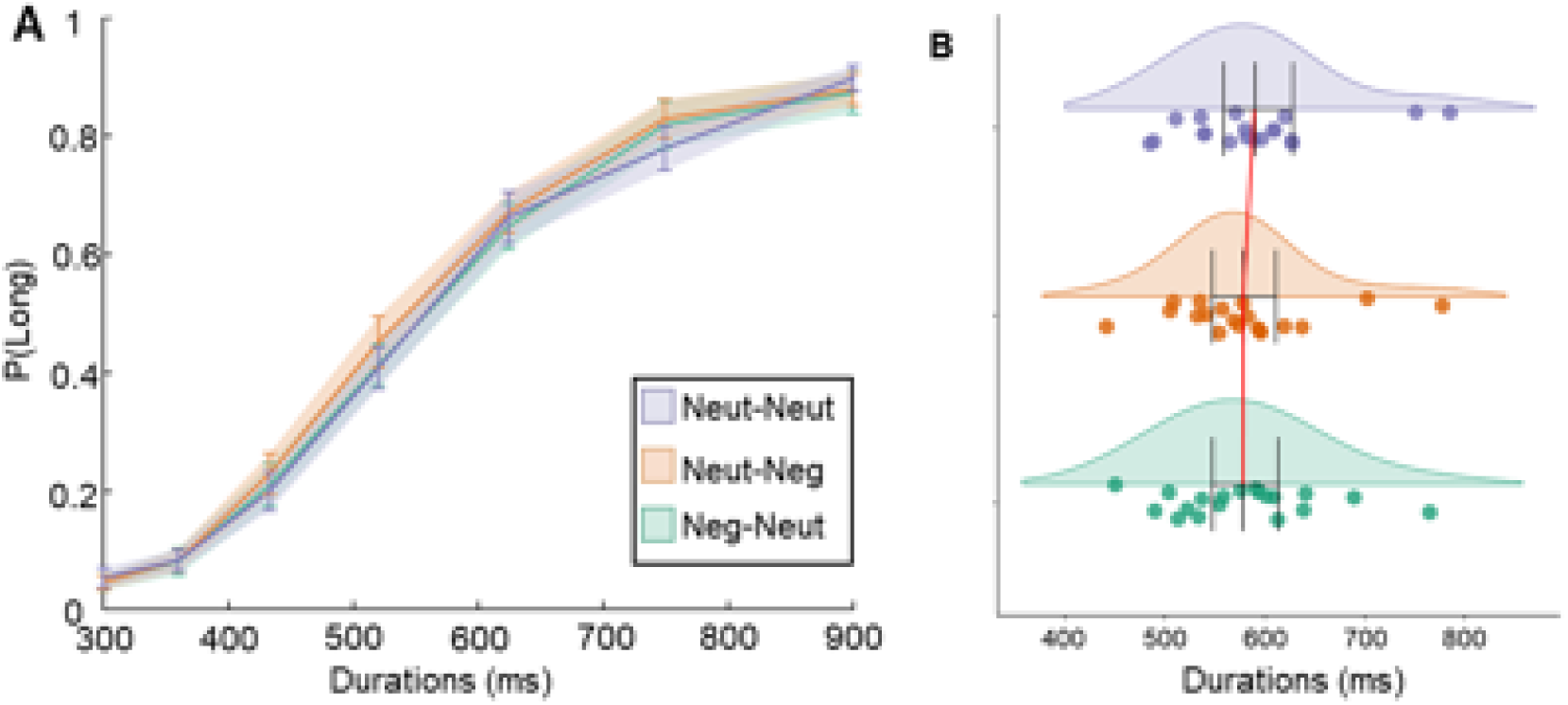
Behavioral results. No significant effect of condition. (A) Psychometric curve displaying the mean proportion of long responses for each duration across condition with standard error bars. (B) Raincloud plot (Allen et al., 2019) displaying bisection points (BP) and 95% confidence intervals for each condition. Slight leftward shift in BP suggests a minor overestimation of time in Neut→Neg and Neg→Neut.

### Electroencephalogram (EEG) effects

### N170

A repeated measures ANOVA of the N170 amplitude at posterior electrodes during face presentation revealed a significant effect of emotion, regardless of presentation order (Emotion: *F*_(1,18)_ = 32.33, *p* <.001, ƞ^2^ = 0.64; Order: *F*_(1,18)_ = 1.16, *p* = 0.30, respectively). Post-hoc Wilcoxon signed-rank tests showed that the N170 amplitude was significantly larger for negative faces then neutral faces regardless of whether participants saw the faces before or after the temporal stimulus (Figure 4; Face1: *W*=178.00, *p* <.001, r_rb_ = 0.87; Face2: *W* = 190.00, *p* <.001, r_rb_ = 1). (Face1-Neg: M = 2.88, SE = 0.50, 95% CI = [1.89, 3.87]; Face1-Neut: M = 1.61, SE = 0.41; 95% CI = [0.82, 2.40]; Face2-Neg: M = 2.28, SE = 0.41, 95% CI = [1.47, 3.09]; Face2-Neut: M = 1.32, SE = 0.32, 95% CI = [0.68, 1.95]).

**Figure 4.**
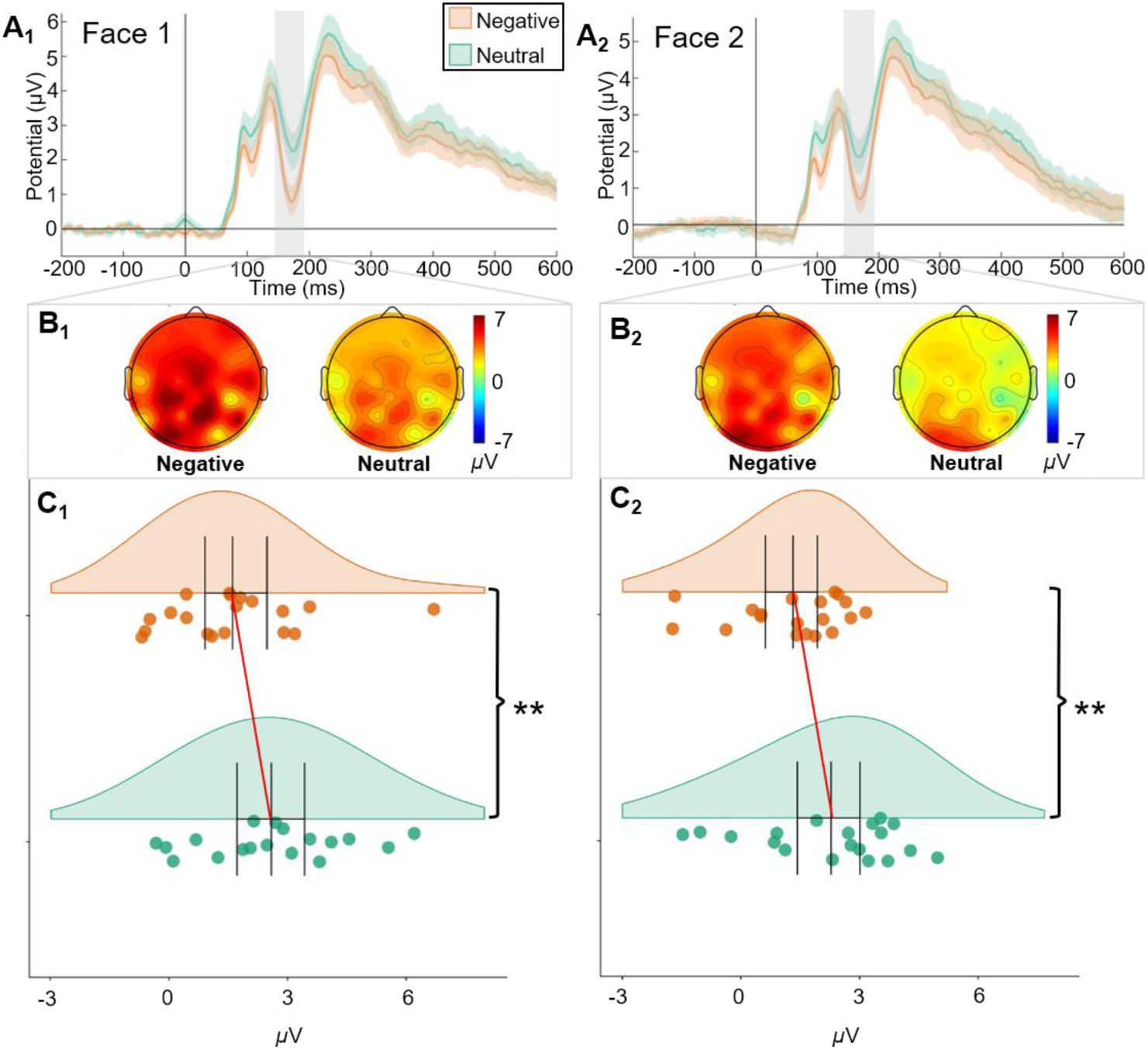
N170 results. (A_1_) First face and (A_2_) Second face ERP waveform analyzed at 145-185ms after face stimuli onset at electrodes P7, P8, PO7, PO8, displaying means and standard errors. (B_1_) First face and (B_2_) second face topography of N170. (C_1_) First face and (C_2_) second face Raincloud plots displaying mean amplitude with confidence intervals. (** = p<.001)

#### N1/CNV

For the N1 and CNV amplitudes (Figure 5), although there was no significant main effect of condition on the N1 (*F*_(2,36)_ = 1.69, *p* = 0.20), there was a marginally significantly difference in the amplitude between the Neut→Neut condition and the Neg→Neut condition (*W* = 50.00, *p* = 0.07) but not between Neut→Neut and Neut→Neg (*W* = 66.00, *p* = 0.26) or Neg→Neut and Neut→Neg (*W* = 87.00, *p* = 0.77). The CNV, on the other hand, had a significant main effect of condition on the amplitude (*F*_(2,36)_ = 3.86, *p* = 0.03, ƞ^2^ = 0.18) with significant difference between each of the three conditions (Neg→Neut/Neut→Neg: *W* = 39.00, *p* =. 02, r_rb_ = −0.59; Neg→Neut/Neut→Neut: *W* = 30.00, *p* =. 007, r_rb_ = −0.68; Neut→Neg/Neut→Neut: *W* = 42.00, *p* = 0.3, r_rb_ = −0.56). (Neut→Neut: M = −3.80, SE = 0.55, 95% CI = [-4.89, −2.71]; Neut→Neg: M = −4.35, SE = 0.62, 95% CI = [-5.55, −314]; Neg→Neut: M = −5.06, SE = 0.60, 95% CI = [-6.24, −3.88]).

**Figure 5.**
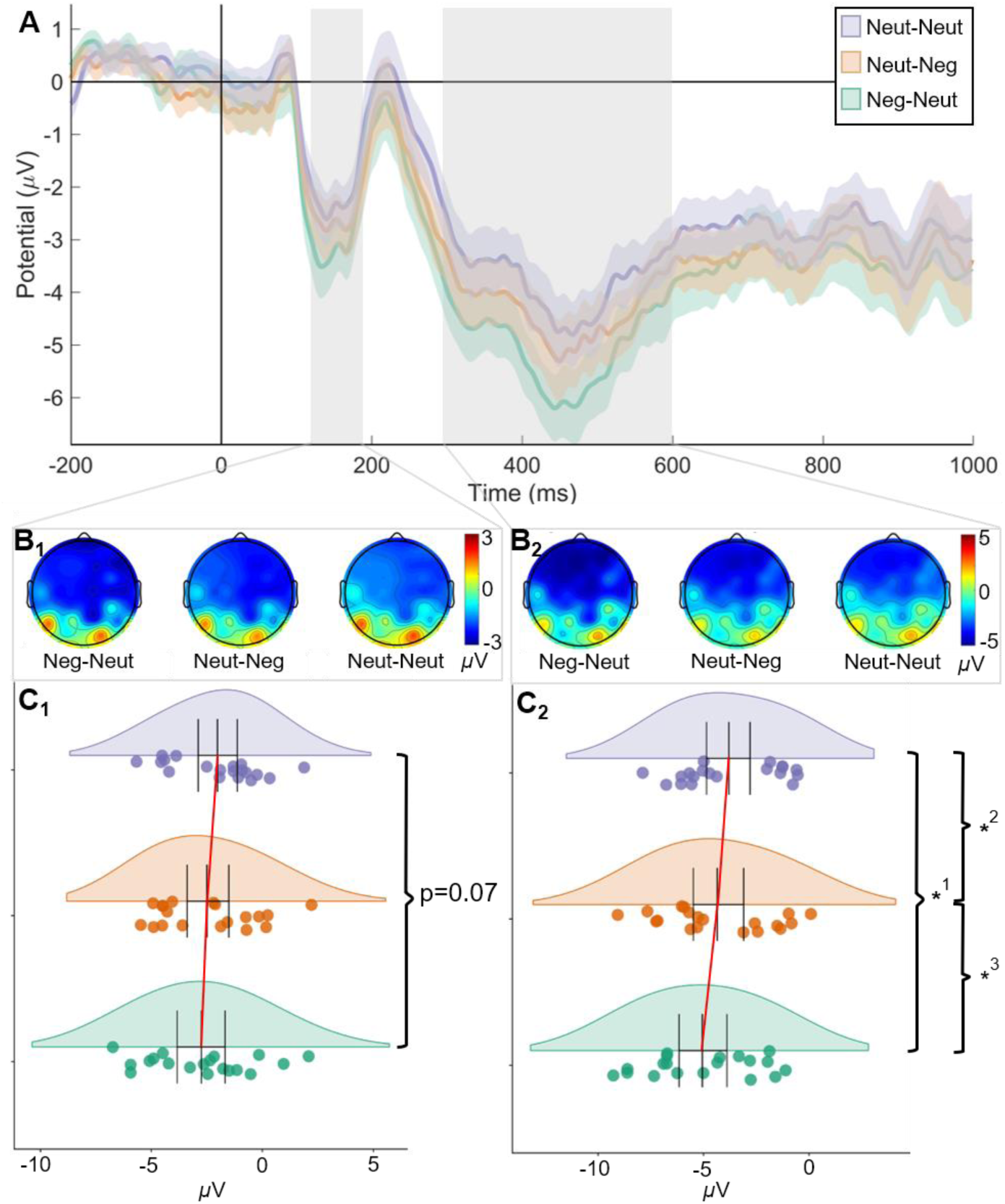
N1/CNV results. (A) N1 and CNV waveform, displaying mean and standard error, analyzed at 150-190ms and 300-600ms after temporal stimulus onset, respectively, at electrodes FCz, Fz, Cz, F1, F2, FC1, FC2, C1, C2. (B_1_) Topography of the N1. (B_2_) Topography of the CNV. (C_1_) Raincloud plots displaying mean amplitude of N1 with confidence intervals. (C_2_) Raincloud plots displaying mean amplitude for CNV with confidence intervals. (p-values: ^*1^ =. 007; ^*2^ =. 032; ^*3^ =. 023)

The significant difference between the Neut→Neg and Neut→Neut conditions in the CNV was an unexpected finding since, at this point in the task, the Neut→Neg and Neut→Neut conditions are identical we did not anticipate there to be a significant difference between them. As an explanation, we considered that participants may have been anticipating another negative face first after experiencing a Neg→Neut trial (Kahn & Aguirre, 2011; Wiener, Thompson & Coslett, 2014). We therefore removed Neut→Neg trials that followed Neg→Neut trials to reduce this potential carryover effect. The repeated measures ANOVA of the filtered data revealed a nearly significant main effect of condition (*F*_(2,36)_ = 2.94, *p* = 0.07, ƞ^2^ = 0.14). Post-hoc analyses revealed that the CNV amplitude for the Neg→Neut remained significantly higher than Neut→Neg (*W* = 42, *p* = 0.03, r_rb_ = −0.56) and Neut→Neut (*W* = 31, *p* = 0.01, r_rb_ = −0.67). Additionally, the Neut→Neg and Neut→Neut CNV amplitudes were no longer significantly different from each other (*W* = 53, *p* = 0.10, r_rb_ = −0.44), consistent with our explanation and original expectations.

#### LPCt

There was no significant effect of condition or response on the LPCT amplitude (*F*_(2,36)_ = 1.49, *p* = 0.24; *F*_(1,18)_ = 1.00, *p* = 0.33, respectively); however, there was an interaction of condition and response (Figure 6; *F*_(2,36)_ = 4.43, *p* = 0.02, ƞ^2^ = 0.20). Specifically, there was a significant difference in the LPCt amplitude for short responses between the Neut→Neg and Neut→Neut condition (*W* = 20.00, *p* =. 001, r_rb_ = −0.79) and a marginally significant difference for short responses between Neg→Neut and Neut→Neg (*W* = 141.00, *p* = 0.07, r_rb_ = 0.48). We also found that the LPCt amplitude for short and long responses were significantly different in the Neut→Neut condition (*W* = 29.00, *p* =. 006, r_rb_ = −0.70). (Neut→Neut_S: M = 7.75, SE = 2.07, 95% CI = [3.68, 11.81]; Neut→Neg_S: M = 5.60, SE = 1.57, 95% CI = [2.52, 8.68]; Neg→Neut_S: M = 7.18, SE = 2.16, 95% CI = [2.94, 11.42]; Neut→Neut_L: M = 6.52, SE = 2.0, 95% CI = [2.59, 10.45]; Neut→Neg_L: M = 6.62, SE = 1.64, 95% CI = [3.40, 9.84]; Neg→Neut_L: M = 6.33, SE = 1.62, 95% CI = [3.16, 9.51]).

**Figure 6.**
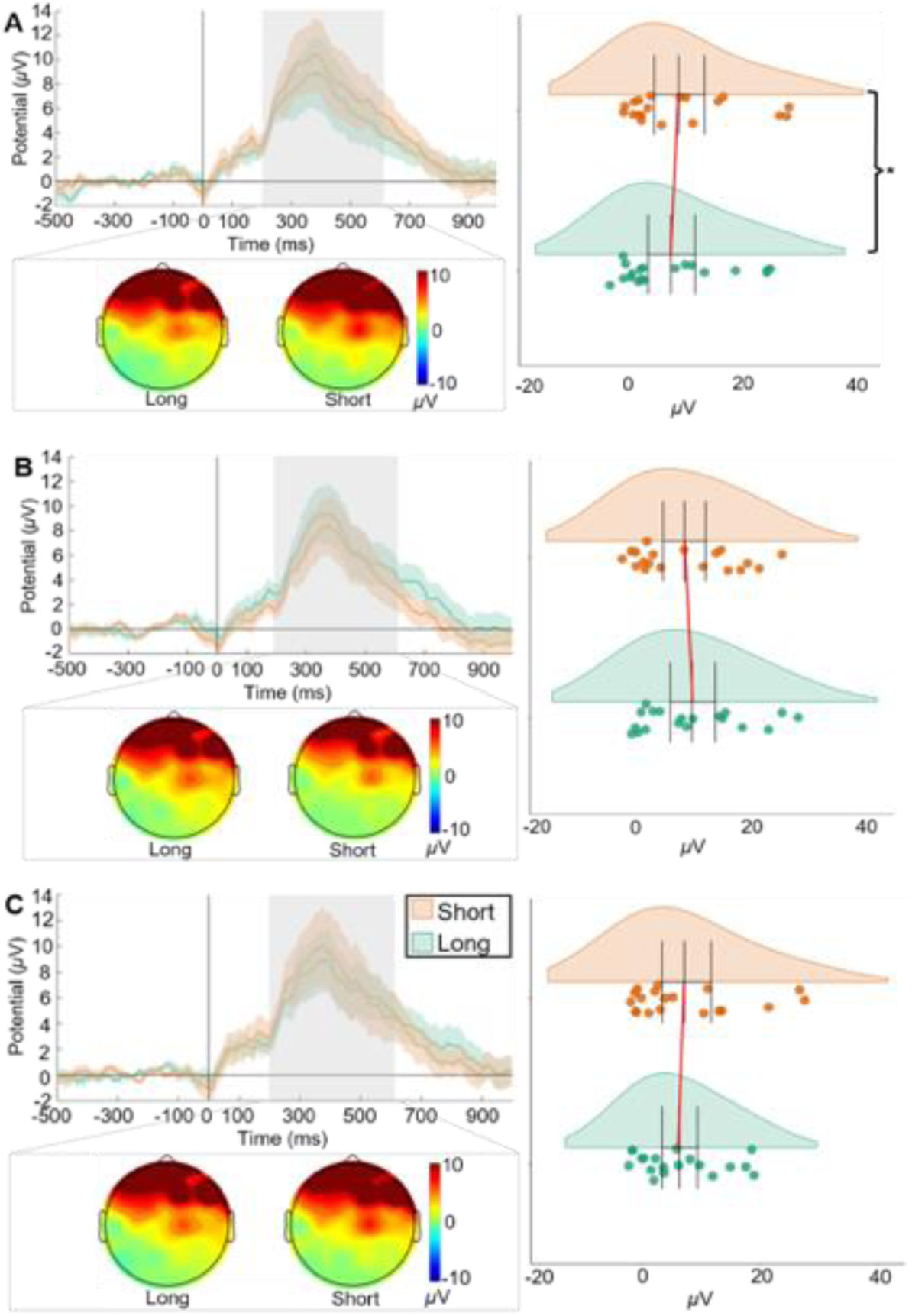
LPCt results. Significant interaction effect, no significant effect of condition or response alone. (A), (B), and (C) ERP waveform showing mean and standard error, topography map, and Raincloud plots showing mean and confidence intervals. Waveform analyzed at 200-600ms after response onset at electrodes FCz, Fz, Cz, F1, F2, FC1, FC2, C1, C2. (A) Neut→Neut condition. Amplitude for ‘Short’ responses significantly more positive than for ‘Long’ responses. (B) Neut→Neg condition and (C) Neg→Neut condition - No significant effect of response. Interaction effects: LPCt amplitude for ‘Short’ responses in the Neut→Neut condition is significantly more positive than in the Neut→Neg condition. Amplitude is near significantly larger in the Neg→Neut condition compared to the Neut→Neut condition. No significant differences in amplitude for ‘Long’ responses between conditions.

## Discussion

In the present study, we found that participants were prone to overestimate time both when an emotional face preceded and, surprisingly, followed a timed stimulus. The CNV results suggest that time was overestimated when the emotional face preceded the temporal stimulus due to a mixture of arousal (as indexed by the face-responsive N170) and attention (as indexed by the timed stimulus N1). Additionally, an interaction effect of condition and response observed on the LPCt suggests that time was also overestimated when the emotional face succeeded the temporal stimulus. This suggests that emotion can affect both the clock and the decision-making stage of the pacemaker-accumulator model.

The CNV signal has previously been associated with the perception of time (Kononowicz, van Rijn, & Meck, 2018). A larger, more negative CNV amplitude has been interpreted to represent a larger number of pulses that have accumulated (Macar & Vidal, 2004; Wiener, et al. 2012); therefore, our results suggest that participants overestimated neutral stimulus duration when preceded by an angry face, as opposed to a neutral face. However, it remains unclear as to whether the emotional impact on the clock stage is specifically due to attention or arousal, as our findings could be interpreted as evidence for both. The face-responsive N170 at posterior electrodes was significantly more negative for angry faces than for neutral faces, suggesting that the angry faces were perceived as more arousing (Hinojosa et al., 2015). However, we also found that the N1 response evoked by the subsequent temporal stimulus was marginally larger when preceded by an angry face, as opposed to a neutral face, suggesting increased attention (Brunia, et al., 2011). Therefore, overestimation could be due to either an increase of the pacemaker rate via arousal, or the gate opening wider via increased attention, or a combination of both. Given previous research and our own findings regarding attentional and arousal effects of emotion on time perception, we suggest that it may not be an either-or process but that both attention and arousal collectively distort time perception in the case of emotional stimuli (Burle & Casini, 2001).

The response locked LPCt revealed that the amplitude varied based on response (short or long) as previously shown by Wiener and Thompson (2015). More notable was an interaction effect between response and condition, with the LPCt amplitude for “short” responses exclusively being significantly lower when participants saw a negative face before responding (Neut→Neg) compared to a neutral face (Neut→Neut). There was also a marginally significant difference in amplitude for “short” responses between Neut→Neut and Neg→Neut suggesting that a negative face before the temporal stimulus still causes some effect at the decision-making stage.

These findings may be related to those that suggest more positive LPCt amplitudes are associated with more difficult discrimination tasks, as evidenced by decreases in accuracy and reaction time associated with higher LPCt amplitudes (Paul et al., 2011). However, if this was the case, we would expect the LPCt amplitude for both short and long responses to increase by similar amounts in the Neut→Neg condition and only slightly in the Neg→Neut condition compared to Neut→Neut. Instead, the LPCt amplitude for short responses decreased in the Neut→Neg condition, compared to Neut→Neut, but remains the same for long responses across all conditions. We therefore suggest that the LPCt results instead reveal a bias in decision-making, specifically against choosing “short” as a response option. This finding could additionally be interpreted as a shift in the so-called “choose-short” effect (Lieving, et al. 2006), in which human exhibit a greater tendency to categorize stimuli as “short” in temporal bisection. Additionally, Wiener & Thompson (2015) also found LPCt amplitudes to more closely match choice probability rather than the timed durations as previously thought. Consequently, we believe that the LPCt amplitude difference for short and long responses in the Neut→Neut condition and the change in LPCt amplitude for only short responses in the Neut→Neg condition and less so in the Neg→Neut condition compared to Neut→Neut represents a bias in decision-making. Therefore, in the control (Neut→Neut) condition, participants are more likely to choose short, evidenced by a larger LPCt for short responses than for long responses, and seeing a negative face prior to responding (Neut→Neg) caused a reduction in this effect.

We note here that the behavioral data did not reveal significant differences between conditions; however, the slight leftward shift in BP for Neg→Neut and Neut→Neg suggests that a negative face before a temporal stimulus and a negative face after the temporal stimulus (before the response) resulted in an overestimation. We do, however, speculate that this weak effect may partially be due to the emotional stimuli chosen and/or the presentation order of the stimuli. Previous research suggests that emotional faces are not as arousing as other emotional stimuli (e.g. IAPS; Britton et al., 2006), therefore, using facial stimuli may have resulted in a weaker effect. Additionally, the emotional stimuli were presented before or after, rather than during, timing or response. This may have also caused a weaker behavioral effect but was necessary in order to analyze important ERPs independently from one another. For example, if the emotional stimuli were displayed during the timing or response phase, it would not be possible to fully differentiate the N170 from the N1 and CNV or the LPCt.

In summary, we found that negative emotional faces not only likely increased the perceived time of an interval but also biased temporal discrimination as evidenced by the CNV and LPCt, respectively. Additionally, our results support that the effect of emotion on time perception may not be exclusively due to arousal or attention alone but rather a combined effect. As far as we are aware, an effect of emotional stimuli on decision bias in time estimation has not been observed, and further no neural evidence to dissociate such an effect. As such, we have also provided more evidence for the LPCt as a measure of decision-making bias with more positive amplitudes relating to choice probability.

